# LIN28 selectively modulates a subclass of let-7 microRNAs

**DOI:** 10.1101/202036

**Authors:** Dmytro Ustianenko, Hua-Sheng Chiu, Sebastien M. Weyn-Vanhentenryck, Pavel Sumazin, Chaolin Zhang

**Author notes:** Equal contribution (Dmytro Ustianenko, Hua-Sheng Chiu). To whom correspondence should be addressed (C.Z.), (P.S.).

## Abstract

LIN28 is a bipartite RNA-binding protein that post-transcriptionally inhibits let-7 microRNAs to regulate development and influence disease states. However, the mechanisms of let-7 suppression remains poorly understood, because LIN28 recognition depends on coordinated targeting by both the zinc knuckle domain (ZKD)—which binds a GGAG-like element in the precursor—and the cold shock domain (CSD), whose binding sites have not been systematically characterized. By leveraging single-nucleotide-resolution mapping of LIN28 binding sites *in vivo*, we determined that the CSD recognizes a (U)GAU motif. This motif partitions the let-7 family into Class I precursors with both CSD and ZKD binding sites and Class II precursors with ZKD but no CSD binding sites. LIN28 *in vivo* recognition—and subsequent 3′ uridylation and degradation—of Class I precursors is more efficient, leading to their stronger suppression in LIN28-activated cells and cancers. Thus, CSD binding sites amplify the effects of the LIN28 activation with potential implication in development and cancer.

## Introduction

MicroRNAs (miRNAs) are a class of small regulatory RNAs of ~22 nt that are involved in essentially all cellular processes. To produce mature miRNAs, the primary transcripts of the miRNA gene (pri-miRNAs) are first cleaved in the nucleus into a hairpin precursors (pre-miRNAs) by the microprocessor complex containing DROSHA and the RNA binding protein (RBP) DGCR8, and then exported to the cytoplasm for further processing by DICER to remove its loop region. One strand of the resulting duplex is incorporated into the RNA-induced silencing complex (RISC) to serve as a template for suppressing target mRNA through complementary base-pairing (Kim et al., 2009).

Let-7 is an ancient family of miRNAs initially discovered as a heterochronic gene in *C. elegans* (Lee et al., 1993) but later found in all bilateral animals (Pasquinelli et al., 2000). In humans, the let-7 family consists of 12 members that are expressed from 8 different loci generated by genomic duplication events during evolution (Hertel et al., 2012). All members of the let-7 family contain an identical seed sequence, the major determinant of target selection, and their targets include oncogenes RAS (Johnson et al., 2005), HMGA2 (Lee and Dutta, 2007; Mayr et al., 2007), c-MYC (Sampson et al., 2007), and multiple genes involved in pluripotency maintenance (Worringer et al., 2014). Interestingly, while the levels of pri- and pre-let-7 are comparable between undifferentiated and differentiated cells, it was reported that mature let-7 are detected only after differentiation of ESCs (Suh et al., 2004; Thomson et al., 2006; Wulczyn et al., 2007), suggesting a post-transcriptional mechanism that suppresses their biogenesis. This suppression was later found to be mediated by an RBP named LIN28 (Heo et al., 2008; Moss et al., 1997; Newman et al., 2008; Rybak et al., 2008; Viswanathan et al., 2008).

The LIN28 protein, consisting of an N-terminal cold shock domain (CSD) and a C-terminal CCHC-type zinc knuckle domain (ZKD), is encoded by two paralogous genes *LIN28A* and *LIN28B* (Figure 1A and Figure S1A). Expression of LIN28 is mainly restricted to ESCs and certain transformed cell lines, but is reactivated in ~15% of tumors (Shyh-Chang and Daley, 2013; Viswanathan et al., 2009). The profound impact of the LIN28/let-7 axis is highlighted by the fact that LIN28 is one of four factors sufficient to reprogram human somatic cells into induced pluripotent stem cells (Hanna et al., 2009; Yu et al., 2007). Consequently, extensive efforts have been made to understand the underlying mechanism of LIN28-mediated let-7 suppression and multiple mechanisms have been proposed. These include blocking of DROSHA processing of pri-let-7 in the nucleus (Newman et al., 2008; Viswanathan et al., 2008); DICER processing of pre-let-7 (Heo et al., 2008; Lightfoot et al., 2011; Rybak et al., 2008); and 3′ end uridylation (Hagan et al., 2009; Heo et al., 2009) which stimulates further degradation of pre-let-7 by the DIS3L2 exonuclease (Chang et al., 2013; Ustianenko et al., 2013).

**Figure 1:**
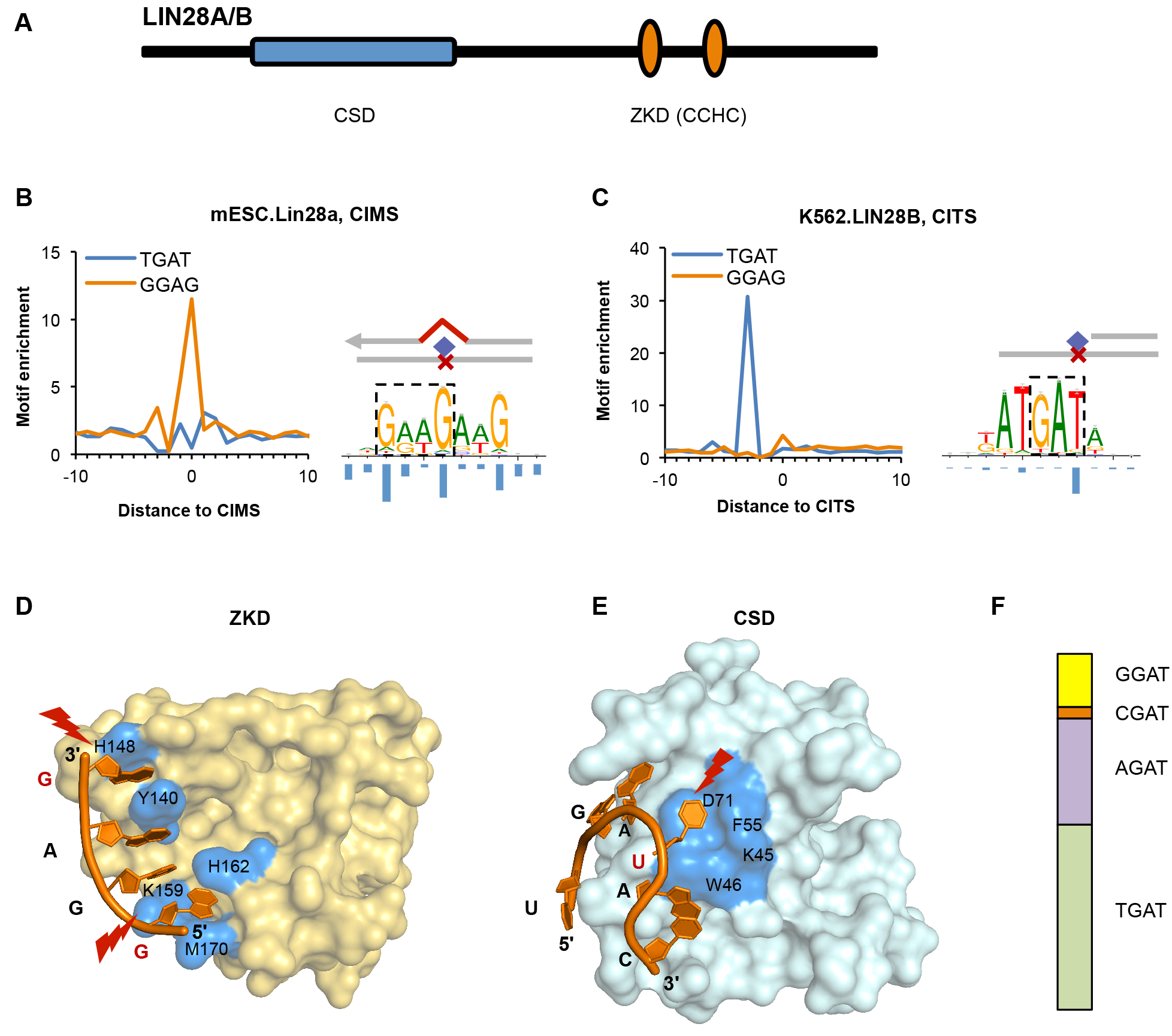
LIN28 cold shock domain (CSD) and zinc knuckle domain (ZKD) recognize distinct sequence motifs as defined by single-nucleotide resolution analysis of CLIP data. Related to Figures S1 and S2. (A) Schematic representation of LIN28 protein domains. (B, C) The ZKD and CSD motifs determined from single-nucleotide resolution analysis of CLIP data. A GGAG motif was identified by modeling sequences around LIN28A CIMS derived from mouse ESCs (B, right panel), and a UGAU motif was determined by modeling sequences around LIN28B CITS derived from K562 cells (C, right panel). The frequency of crosslinking at each motif position is shown under the motif logos. The enrichment of GGAG and UGAU tetramers around CIMS or CITS is shown on the left of each panel. (D, E) The X-ray crystallographical structure of LIN28A ZKD (D) and CSD (E) in complex with let-7g hairpin (PDB accession: 3TS2). Residues that are in direct contact with RNA are highlighted in blue. The crosslinked nucleotides are indicated in red and highlighted. (F) Frequency of tetramers conforming to the NGAU consensus in LIN28B eCLIP data from K562 cells.

Both the CSD and ZKD are involved in recognition of the pre-let-7 through the terminal stem-loop structure, as demonstrated by extensive mutational analysis, *in vitro* miRNA processing assays, and LIN28/pre-let-7 cocrystal structure (Heo et al., 2009; Mayr et al., 2012; Nam et al., 2011; Piskounova et al., 2008). It has been well established that the ZKD recognizes a GGAG-like motif located in the stem loop structure. In human, this motif is present in all members but one of the let-7 family (Triboulet et al., 2015), and it is crucial for stabilizing the LIN28 and pre-let-7 complex and recruiting the terminal uridine transferase (TUTase) that uridylates pre-let-7 (Wang et al., 2017).

Multiple studies have reported that the CSD has a higher affinity to several tested pre-let-7 members than the ZKD (Mayr et al., 2012; Nam et al., 2011; Wang et al., 2017), but its sequence specificity is under debate. Analysis of the LIN28/pre-let-7 co-crystal structure revealed that the CSD interacts with the single stranded loop area of pre-let-7 hairpin and is predicted to have a preference for the GNGAY sequence (Y=pyrimidine; N=any base). However, due to variations in the loop region among the 12 let-7 family members, this motif is only present in a subset of pre-let-7. Assuming all pre-let-7 family members are uniformly suppressed by LIN28, Nam et al. proposed that the CSD has weaker sequence specificity, so that it can adopt to different substrate sequences (Nam et al., 2011). The CSD was also reported to have a preference for pyrimidine rich sequences and it was suggested that its interaction with the loop region of pre-let-7 might induce a conformational change that exposes the GGAG motif in the hairpin (Mayr et al., 2012).

In addition to let-7 miRNAs, several recent studies using crosslinking and immunoprecipitation followed by high-throughput sequencing (HITS-CLIP or CLIP-seq) demonstrated that LIN28 recognizes thousands of mRNA transcripts, and might play a role in regulating RNA splicing and translation through less characterized mechanisms (Cho et al., 2012; Graf et al., 2013; Hafner et al., 2013; Wilbert et al., 2012). Analysis of LIN28 binding sites in mRNA revealed an enrichment of GGAG-like sequences corresponding to the ZKD binding motif. However, these studies have so far provided limited insights into the sequence specificity of the CSD and its contribution to *in vivo* LIN28-RNA interactions, possibility due to insufficient resolution for deconvoluting the bipartite LIN28 binding motif.

In this study, we characterized the *in vivo* binding specificity of LIN28 using single-nucleotide resolution maps of thousands of LIN28 binding sites in mRNA derived from CLIP data. Our analysis confirmed the GGAG motif is recognized by the ZKD. Importantly, we identified a novel, high-confidence CSD binding motif— (U)GAU—which is reminiscent of the CSD-binding consensus sequence proposed based on the LIN28/pre-let7 co-crystal structure (Nam et al., 2011). We further observed that LIN28 binds much more robustly to the subclass of pre-let-7 harboring (U)GAU (Class I), but not the other subclass without the motif (Class II), both in mouse ESCs and human cancer cell lines. Consequently, Class I let-7 family members are efficiently uridylated and suppressed *in vivo*, while the impact of LIN28 on Class II let-7 family members is much more moderate. The selective suppression of the two subclasses of let-7 was also observed in multiple tumor types where LIN28 expression is reactivated, implying a potential role of this selective suppression model in tumorigenesis.

## Results

### Identification of a novel CSD binding motif from single-nucleotide resolution analysis

Given the bipartite nature of LIN28 RNA-binding domains, we postulated that a single-nucleotide-resolution map of LIN28-RNA interaction sites would help to better characterize its binding specificity. To this end, we took advantage of the computational approaches previously developed in our laboratory to infer the precise protein-RNA crosslink sites from CLIP data by identifying crosslink-induced mutation sites (CIMS) and truncation sites (CITS) (Weyn-Vanhentenryck et al., 2014; Zhang and Darnell, 2011). We applied this method to two in-depth LIN28 CLIP datasets: LIN28A HITS-CLIP performed in mouse ESCs (Cho et al., 2012), and LIN28B CLIP derived from two human cell lines K562 and HepG2 using a modified CLIP protocol named eCLIP (Van Nostrand et al., 2016) (for this study, we mainly describe results from K562 cells, as the results obtained from HepG2 cells are very similar). Due to the different protocols used to generate these CLIP libraries, we expected HITS-CLIP to capture only CIMS and eCLIP to be enriched in CITS (see Discussion). Following our established pipeline (Shah et al., 2017), we identified 50,292 substitution CIMS from LIN28A HITS-CLIP data and 22,673 CITS from LIN28B eCLIP data in K562 cells. Consistent with the previous analysis (Cho et al., 2012), we observed a striking enrichment of the GGAG motif at CIMS inferred from HITS-CLIP data (11.5 fold at position 0), indicating the predominant crosslinking of the first G of the GGAG motif (Figure 1B left panel and Figure 1D). Surprisingly, we found only very moderate enrichment of the GGAG motif around CITS inferred from eCLIP data (4.2 fold at position 0 as compared to ~2 fold in neighboring positions; Figure 1C left panel), suggesting a possibility that these binding sites reflect a second mode of LIN28-RNA interaction.

To better understand the binding specificity of LIN28, we performed *de novo* motif analysis using an algorithm we developed to simultaneously model the binding specificity of an RBP and its crosslinking position in the sequence motif (see Methods). This algorithm recovered the GGAG motif from sequences around CIMS, with more degeneracy allowed between the first and last guanines, which is consistent with previous structural and mutational analyses (Loughlin et al., 2011; Nam et al., 2011). Intriguingly, applying this method to sequences around CITS revealed a distinct AUGAU motif, with predominant crosslinking in the last uridine. The tetramer motif UGAU is strikingly enriched in sequences around CITS (31 fold at position -3, corresponding to crosslinking to the last uridine), but not CIMS (Figure 1B and C), suggesting its potential importance for LIN28 binding to thousands of mRNA transcripts.

After careful examination, we noticed that the UGAU motif largely resembles the CSD-binding consensus GNGAY proposed from X-ray crystal structural analysis of LIN28 in complex with pre-let-7d, pre-let-7f-1 and pre-let-7g (Figure 1E). The position of the crosslink sites in this newly identified motif is highly consistent with the RNA contact of the LIN28 CSD (Figure 1E). Presence of the purines in the middle of motif (U<u>GA</u>U) is critical for reaching the protein surface while the last pyrimidine is essential due to the steric hindrance that is imposed by the surrounding amino acids (Nam et al., 2011). Given that a uridine before GAU does not seem to be crucial for LIN28 binding to pre-let-7 (Nam et al., 2011), we examined the other variants of the GAU motif in LIN28 binding sites in mRNAs. Indeed, we observed UGAU, AGAU, GGAU and AGAU are all enriched around CITS to a varying degree (Figure 1F), suggesting that a GAU core motif is the primary determinant of CSD binding.

To confirm the involvement of the newly identified CSD motif in the bipartite binding of LIN28, we examined the enrichment of both GGAG and GAU motifs in sequences around the most robust LIN28 CLIP tag peaks in mRNAs independent of the identified crosslink sites. Both motifs were found enriched in HITS-CLIP as well as in eCLIP data sets, with GAU motif being highly represented 5-30 nucleotide upstream of the GGAG motif (Figure S1B-D). We also predicted LIN28 binding sites in mRNA transcripts by using our mCarts algorithm to identify likely functional clusters of conserved LIN28 motif sites (Weyn-Vanhentenryck and Zhang, 2016; Zhang et al., 2013). The clusters predicted with conserved GAU motifs better overlap the LIN28 CLIP data than those predicted with GGAG. Critically, the best performance was achieved by a hybrid model that allows any combination of GAU and GGAU motifs (Figure S2). These results indicate that the presence of both binding elements contributes to high-affinity interaction of LIN28 and mRNA targets, which is consistent with the bipartite mode of LIN28 - pre-let-7 interaction. Together, our analysis suggests the sequence-specificity of both LIN28 CSD and ZKD and the importance of the bipartite binding motif for *in vivo* protein-RNA interaction.

### Selective recognition of pre-let-7 is modulated by the CSD binding site

Since let-7 pre-miRNAs are the best known targets of LIN28, we investigated whether the newly identified CSD binding motif fits the *in vivo* recognition pattern of LIN28 to let-7 precursors. Examination of pre-let-7 sequences suggests that the whole family can be divided into two subclasses based on the presence of the CSD binding motif (Figure 2A). Class I let-7 family members contain both GAU and GGAG-like motifs (CSD+). These include pre-let-7b, pre-let-7d, pre-let-f-1, pre-let-7g, and mir-98. We also include pre-let-7i in Class I, which has GAC, a variant of GAU predicted to be compatible with the CSD structure (Nam et al., 2011); in this case the uridine before GAC also matches the CSD binding consensus we determined. Class II let-7 family members have no GAU motif in the terminal loop region of pre-miRNA (CSD-). This subclass includes pre-let-7a-1/2/3, pre-let-7c, pre-let-7e, and pre-let-7f-2. All Class II members but one (pre-let-7a-3) have the GGAG-like motif. Interestingly, it was previously reported that pre-let-7a-3 completely escapes LIN28-mediated suppression (Triboulet et al., 2015). However, the distinction of Class I and Class II let-7 family members with respect to LIN28 binding and LIN28-mediated suppression is unclear.

We hypothesized that if the (U)GAU motif uncovered from analysis of tens of thousands of LIN28 mRNA binding sites reflects the *in vivo* binding specificity of LIN28 CSD, its presence should also be important for LIN28 recognition of let-7 precursors. Since multiple studies consistently reported higher binding affinity of the CSD to pre-let-7 compared to the ZKD (Mayr et al., 2012; Nam et al., 2011; Wang et al., 2017), we predicted that LIN28 binds Class I pre-let-7 much more robustly than Class II pre-let-7 *in vivo.* To validate this prediction, we examined LIN28 binding to all pre-let-7 family members identified by CLIP data. Intriguingly, while robust LIN28B CLIP tag clusters were found for all of the Class I pre-let-7’s, very few CLIP tags were detected from Class II let-7’s in K562 and HepG2 cells. The difference of the two classes is statistically significant after controlling the abundance of pre-miRNA expression (P=0.011, ANOVA; Figure 2B). The distinction can be most clearly observed in let-7 family members expressed from poly-cistronic locus as a single primary transcript (e.g., pre-let-7d and pre-let-7f-1 in Class I versus pre-let-7a-1 in Class II, Figure 2D. See additional examples in Figure S3) (Wang et al., 2011). Moreover, very similar observations were made from LIN28A HITS-CLIP in mouse ESCs (Figure 2E and Figure S3B), suggesting that the selective binding of LIN28 to Class I versus Class II let-7 precursors modulated by the CSD is not specific for LIN28A or LIN28B, or the cellular contexts we examined.

**Figure 2:**
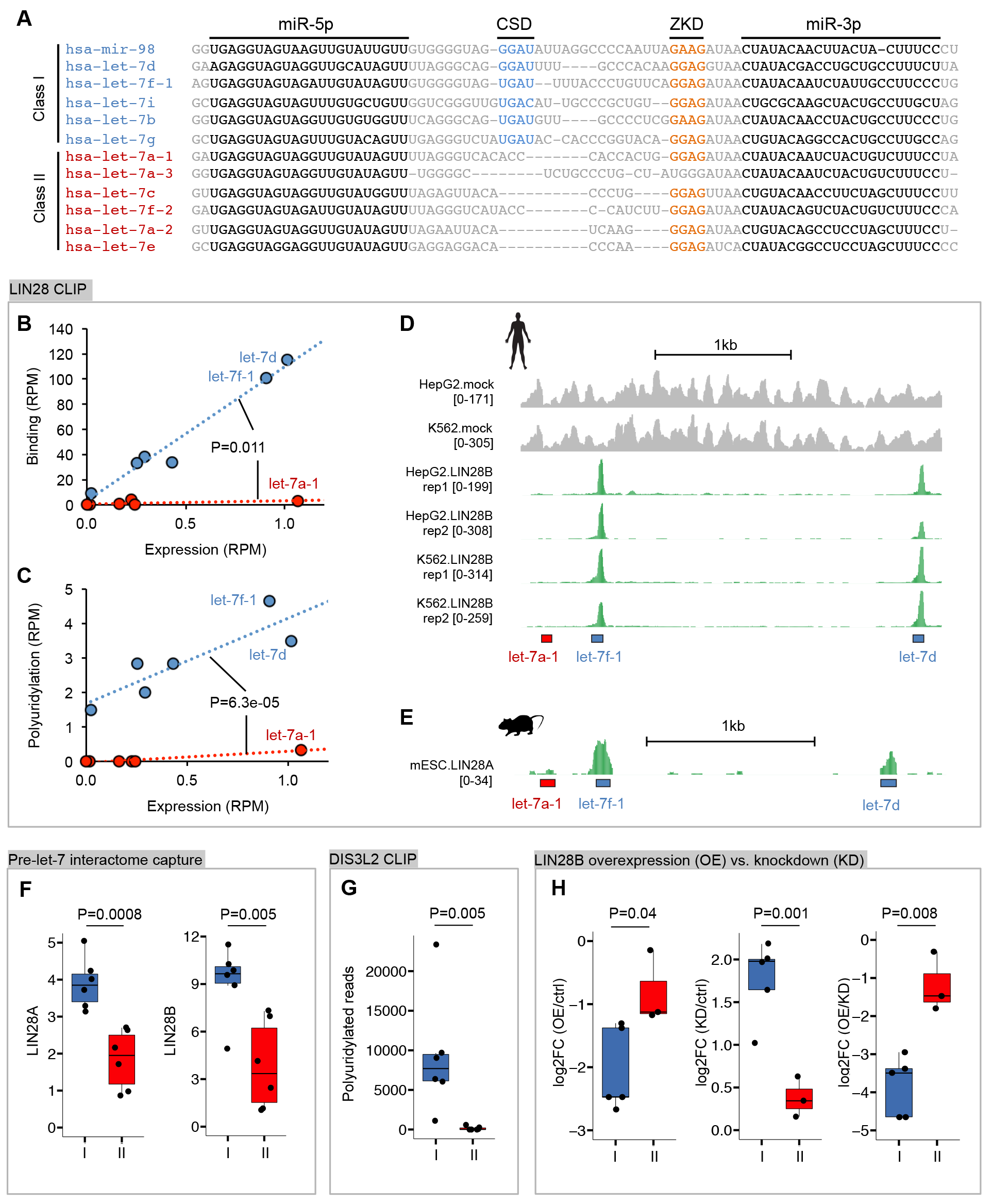
The CSD modulates LIN28 binding on class I let-7 precurosrs, and their 3′ uridylation and degradation. Related to Figure S3. (A) Multiple sequence alignments of pre-let-7 hairpins. Sequences corresponding to mature miRNAs, and binding sites of LIN28 CSD and ZKD are indicated. Let-7 family members are divided into two sub-classes, denoted Class I and Class II, depending on the presence of the CSD motif GAU (GAC in the case of let-7i). (B, C) Quantification of LIN28B binding and 3′ uridylation on let-7 pre-miRNAs in K562 cells. In each panel, the y-axis shows the total number of unique CLIP tags (B), or the number of poly-uridylated CLIP tags (C), expressed in reads per million (RPM), that overlap with each pre-let-7. The x-axis shows the number of mock CLIP tags (input) expressed in RPM reflecting the abundance of the pre-let-7. The difference between LIN28 binding or uridylation after controlling for pre-miRNA abundance is tested using ANOVA. (D, E) LIN28 binding to different let-7 family members in human HepG2 and K562 cells (D) and mESCs (E) using the let-7a-1/7f-1/7d poly-cistronic miRNA locus as an example. The number of mock (gray) and IP (green) tags in each genomic position is shown, and the positions of the pre-miRNA hairpins are indicated at the bottom. (F) LIN28A/B RNA-mediated protein pull-down using different pre-let-7 family members as a bait. The normalized count of mass spectrometry-identified peptides is shown for each protein and each bait (Class I and Class II pre-let-7). The boxplots indicate the interquartile range of each subclass. The difference between the two subclasses was evaluated by a t-test. (G) Boxplot showing the level of uridylation for the two subclasses of let-7 family members from DIS3L2 CLIP in HEK293 cells. Wilcox rank sum test was used to evaluate the difference between the two subclasses. (H) Changes in the expression of mature let-7 miRNA upon perturbation of LIN28B levels (overexpression or knockdown) in HEK293 cells. The difference between the two subclasses was evaluated by a t-test.

To further validate our hypothesis and exclude the possibility that the selective binding observed from CLIP data is due to technical bias (e.g., differences in crosslinking efficiency), we examined a recently published dataset of pre-miRNA-binding interactomes in 11 cell lines, in which pre-miRNA-interacting proteins were captured using an RNA-mediated protein pull-down assay followed by mass spectrometry analysis (Treiber et al., 2017). A number of miRNA precursors including all 12 members of the let-7 family were used as a bait to identify protein factors that specifically recognize the pre-miRNA fold. Importantly, equal amounts of pre-miRNA oligo were used as a bait, so that the amount of protein pull down directly reflects the binding affinity. We compared the number of LIN28 mass spectrometry-identified peptides between Class I and Class II pre-let-7 family members. Both LIN28A and LIN28B showed a greater preference for binding of Class I let-7 (P=0.0008 and 0.005, respectively, t-test; Figure 2F), providing direct biochemical support for selective recognition of let-7 family members by LIN28.

### Class I let-7 miRNAs are selectively uridylated and suppressed by LIN28

The major functional outcome of the LIN28 and pre-let-7 interaction is suppression of mature miRNA levels. One important mechanism of the suppression is LIN28-mediated recruitment of TUT4 polymerase, which modifies the 3′ end of pre-miRNA with a stretch of uridines, and stimulates degradation of pre-miRNA by DIS3L2 exonuclease (Chang et al., 2013; Ustianenko et al., 2013). To evaluate whether selective binding of LIN28 to Class I versus Class II let-7 precursors has any impact on their suppression through the TUT4/DIS3L2 pathway, we referred to a previously published DIS3L2 CLIP analysis (Ustianenko et al., 2016). In this study, a catalytically inactive mutant of DIS3L2 exonuclease with intact RNA binding abilities was used to identify a variety of uridylated RNA transcripts in HEK293 cells. We compared the number of uridylated pre-let-7 DIS3L2 CLIP tags between the two let-7 subclasses and found that Class I pre-let-7’s exhibit up to 20-fold greater uridylation levels compared to Class II pre-let-7’s (P=0.005, Wilcox rank sum test; Figure 2G). Similarly, we compared uridylation levels of let-7 precursors detected in the LIN28B eCLIP data, and found that Class I precursors exhibit significantly higher uridylation levels compared to Class II precursors, regardless of their expression levels (P=6.3e-5, ANOVA; Figure 2C). These observations confirmed that the high affinity LIN28 binding in Class I pre-let-7 mediated by both CSD and ZKD is required for their efficient uridylation *in vivo.*

As 3′ uridylated pre-let-7 is expected to be degraded by DIS3L2, we directly examined whether LIN28 selectively suppresses Class I versus Class II let-7 family members. To this end, we examined the abundance of the individual let-7 family members upon manipulation of LIN28 protein levels either by overexpression or siRNA-mediated knockdown in HEK293 cells (Hafner et al., 2013). We found that overexpression of LIN28B resulted in stronger repression of Class I let-7 compared to Class II let-7; conversely, knockdown of LIN28B resulted in more derepression of Class I let-7 compared to Class II let-7 (P<0.05 in all comparison, t-test; Figure 2H). The same pattern was observed in additional datasets derived from similar experiments (Powers et al., 2016; Wilbert et al., 2012), although the distinction between Class I and Class II let-7 family members was not discussed in the original studies (see Discussion). These observations confirmed that the CSD binding site in Class I let-7 family members plays an important role in determining the efficiency of LIN28-dependent suppression of miRNA biogenesis *in vivo.*

### Class I let-7 miRNAs are selectively suppressed following LIN28 reactivation in human cancers

LIN28 activation, followed by loss of let-7, is a hallmark of cancer etiology (Balzeau et al., 2017). To investigate the suppression of let-7 microRNAs by LIN28 in the context of tumorigenesis, we performed a pancancer analysis of fourteen tumor types for which both mRNA and microRNA expression was profiled by The Cancer Genome Atlas (TCGA) using deep sequencing. In total, we found that LIN28B and LIN28A are variably expressed (mean absolute deviation greater than zero) in six and two tumor types, respectively. In each of these tumor types, the expression of let-7 miRNAs was significantly anti-correlated with that of LIN28 (Figure S4), suggesting suppression of let-7 following LIN28 reactivation, which is consistent with previous studies (Viswanathan et al., 2009). Importantly, in these contexts, Class I and II miRNAs demonstrate variable response to LIN28 activation with Class I miRNAs showing significantly stronger anti-correlation with both LIN28B (Figure 3A) and LIN28A (Figure S5A) expression. Furthermore, the difference was most evident in samples with high LIN28 abundance (Figure 3B and Figure S5B). Taken together, our results suggested that Class I miRNAs are selectively suppressed following LIN28 reactivation in cancer, and that CSD binding sites serve to amplify the LIN28 regulation of Class I miRNAs.

**Figure 3:**
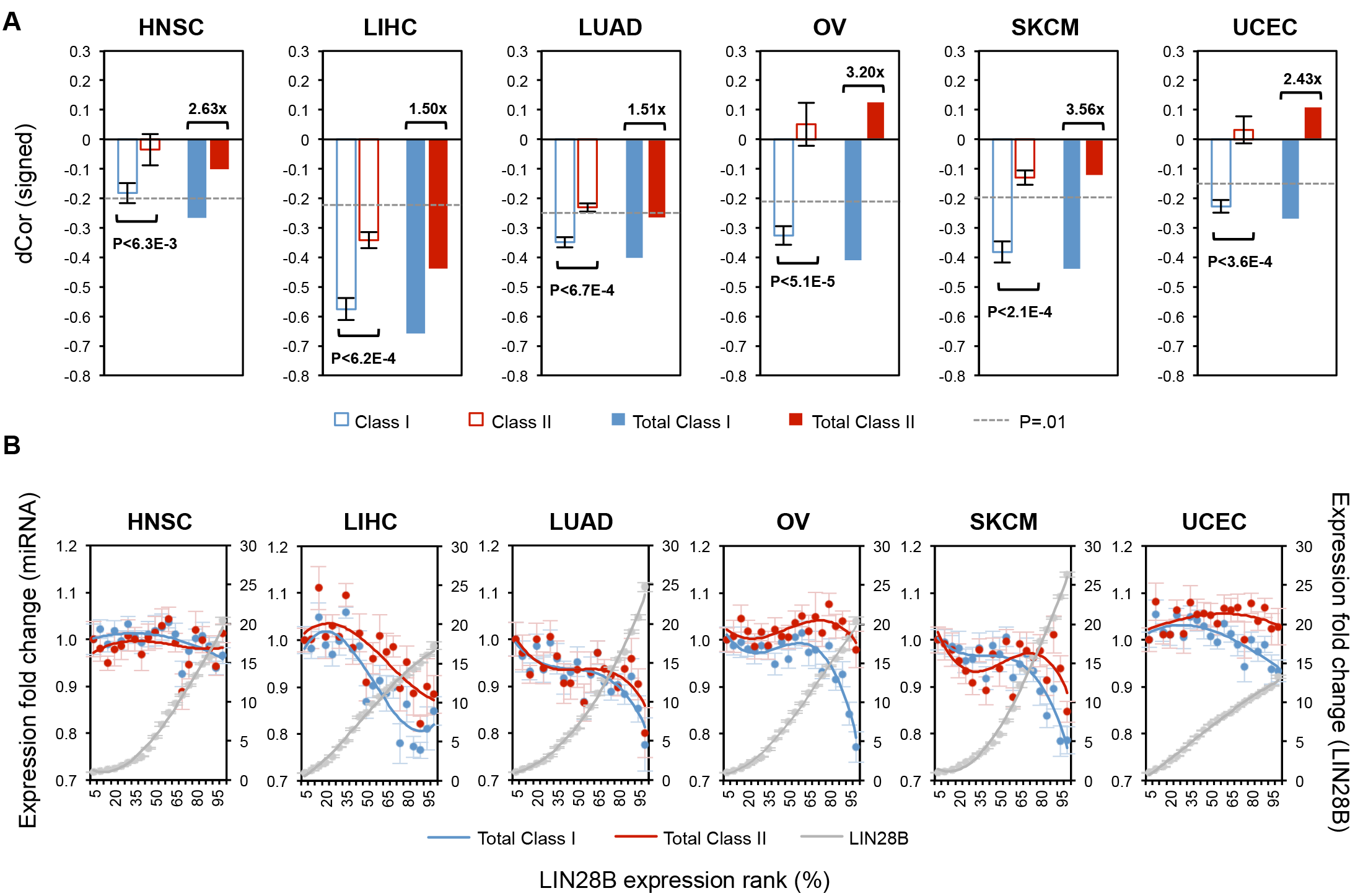
Selective suppression of Class I let-7 in tumor samples with LIN28B reactivation. Related to Figures S4 and S5. (A) Class I miRNAs showed stronger downregulation by LIN28B than Class II miRNAs. For each tumor type, average distance correlation (dCor) estimated between LIN28B and miRNAs from each class are given on the left (hollow bars); the sign is designated by Spearman’s correlation, and p-values estimated by Mann-Whitney U test. Error bars represent standard error of the mean (S.E.M.). Pooled reads across miRNA classes produced total expression per class and their dCor with LIN28B expression is given on the right (solid bars) for each tumor type; the p<0.01 cutoff, estimated by permutation testing, is given in broken gray lines. (B) The response of Class I and II miRNAs to changes in LIN28B expression. Samples were binned into 20 same-size bins according to LIN28B expression. Each bin is represented by the average fold change of total expression in each class relative to the first bin across samples in the bin, and curves were fit to a polynomial distribution with order 3. Similarly, LIN28B average expression fold changes are given on the right axis; S.E.M. are shown.

## Discussion

Due to the important role of the LIN28/let-7 axis in developmental biology and cancer, the mechanisms underlying post-transcriptional suppression of let-7 miRNAs by LIN28 have been a subject of intensive investigation. These studies have revealed the fascinating degree of complexity, which is derived, at least in part, from the plasticity of protein-RNA interactions. In general, each RNA-binding domain of an RBP recognizes a short and degenerate sequence or structural motif. Therefore, specificity has to be achieved through combinations of multiple domains, which allow an expansion of the RNA pool that is regulated in both sequence and structure-dependent manners (Lunde et al., 2007). In the case of LIN28, bipartite binding sites recognized by the CSD and ZKD are required for high-affinity interactions of LIN28 with substrate RNAs, including let-7 pre-miRNAs.

It has been widely believed that all mammalian let-7 miRNAs, with the exception of hsa-let-7a-3 or its homolog, are suppressed by LIN28 through a similar mechanism (Triboulet et al., 2015). This model postulates that the major determinant of specificity is the GGAG-like RNA element recognized by the ZKD while the CSD contacts the terminal loop of pre-let-7, which varies among different let-7 precursors, with limited specificity but higher affinity. To target let-7, LIN28 initially scans RNA through the CSD until the ZKD locates the GGAG element, which stabilizes the LIN28 binding (Nam et al., 2011; Wang et al., 2017), and either blocks microprocessors such as DROSHA and DICER or recruits downstream effectors such as TUTase4/7 that trigger 3′ uridylation and degradation of pre-let-7. As a result, previous reports that investigated the general mechanisms of LIN28/let-7 interaction and LIN28-dependent let-7 biogenesis frequently tested only one or a few selected miRNAs without distinguishing between different let-7 family members. However, examination and comparison of the results from multiple studies suggests that the impact of LIN28 varies across the let-7 family members. While some of these seemingly conflicting results could be due to variability of cellular contexts and experiments, we conjectured that they might also reflect unknown mechanisms that cannot be accounted for by the uniform suppression model.

The importance of the GGAG-like element (e.g., GGAG, GGUG and GAAG) for recognition by LIN28 ZKD is well established (Heo et al., 2009; Mayr et al., 2012). In addition, this interaction was demonstrated to be essential and sufficient for *in vitro* uridylation of let-7g and let-7f-1 (Faehnle et al., 2017; Wang et al., 2017). However, given the relatively weak binding affinity of the ZKD and steric conformation of pre-miRNA, the GGAG motif alone is likely insufficient for high-affinity LIN28 binding and subsequent let-7 suppression *in vivo* (Nam et al., 2011; Wang et al., 2017); efficient suppression of let-7 by LIN28 thus has to rely on the interaction of the CSD with the terminal loop region of pre-let-7. Given the substantial divergence of the terminal loop sequence among let-7 family members, the assumption that LIN28 interacts with all members with similar binding affinity to uniformly suppress let-7 expression might be arguable. However, this question cannot be answered without a precise understanding of the LIN28 CSD binding specificity.

Our single-nucleotide-resolution analysis of tens of thousands of LIN28 binding sites in mRNA using recent eCLIP data unexpectedly uncovered a novel motif (U)GAU. A similar motif GNGAY was proposed to be the consensus binding site of the CSD from the LIN28/let-7 crystal structures (Nam et al., 2011). However, the previous prediction was based on a very limited number of sequences (i.e., pre-let-7d, pre-let-7f-1 and pre-let-7g, all of which contain (U)GAU, for which structures were determined), making it unclear whether this consensus precisely reflects the specificity of the CSD. Partial representation of the motif among let-7 family members is also inconsistent with the uniform suppression model, as well as other studies reporting conflict results of the CSD binding specificity (Mayr et al., 2012). Therefore, without additional support for the significance of the GNGAY consensus, it was postulated that the binding specificity of the CSD is limited, allowing it to adopt to other variable sequences found in all let-7 family members (Nam et al., 2011; Wang et al., 2017).

Our confidence in the (U)GAU motif requirement for high-affinity LIN28 interaction was initially based on its striking enrichment in tens of thousands of LIN28 binding mRNA targets. So why does the eCLIP data capture a distinct LIN28 motif compared to previous CLIP analyses which only identified a GGAG motif (Cho et al., 2012; Graf et al., 2013; Wilbert et al., 2012)? While speculative, this discrepancy is probably due to differences in the protocols used to prepare CLIP libraries. All previous studies using LIN28 CLIP cloned immunoprecipitated RNA crosslinked to LIN28 through ligation of 3′ and 5′ RNA linkers, followed by reverse transcription and PCR amplification using primers that matches the linker sequences. Due to the irreversibility of the crosslinking, it was demonstrated that the residual amino acid-RNA adducts can interfere with reverse transcriptase, sometimes resulting in premature truncation of the cDNA. Only read-through CLIP tags including a subset carrying crosslink-inducted mutations (Zhang and Darnell, 2011) were captured by these protocols, while truncated tags were lost during PCR amplification. On the other hand, eCLIP, among several other similar protocols (Konig et al., 2010), are able to capture both truncated and read-through tags using different cloning strategies. Whether RT enzyme predominantly stops at or reads through crosslink sites depends on the identity of the amino acid-RNA adducts as well as properties of the RT enzyme and other experimental conditions (Van Nostrand et al., 2017). For example, we previously demonstrated that another RBP, RBFOX, can be crosslinked to its binding element UGCAUG at either G2 or G6, which results in predominant read-through and premature stop, respectively. If both CSD and ZKD can be crosslinked to different positions in the bipartite binding site at which they directly contact, the two crosslink sites could also affect RT very differently. For example, crosslinking of the GGAG motif with the ZKD could result in frequent read-through detected in earlier CLIP assays, while crosslinking of the (U)GAU with the CSD could predominantly result in truncations that can only be detected by eCLIP and other improved CLIP protocols. Interestingly, a recent study investigated the crosslinking between LIN28 and pre-let-7f in an *in vitro* binding assay (Ransey et al., 2017). This study confirmed the crosslinking at the guanines in the GGAG motif by CIMS analysis, but also found a predominant crosslink site at the last uridine of the GAU element by tandem mass-spectrometry, which is consistent with our results from genome-wide analysis of *in vivo* LIN28 binding sites.

As the CSD-binding motif is important for LIN28 high affinity interaction with pre-let-7 and the refined motif is not found in all pre-let-7 family members, its presence would divide the let-7 miRNA family into two subclasses. Only the subclass that possesses both GAU and GGAG-like elements (Class I) is predicted to be efficiently targeted by LIN28, implying that the current uniform suppression model has to be modified. We thus propose a new selective suppression model where the presence of the CSD binding element in Class I let-7 family members modulates the efficiency of LIN28-dependent suppression (Figure 4). This model has found strong support from multiple lines of evidence using datasets independently generated by different laboratories. This model fits well with our observation that Class I let-7 family members show much stronger *in vivo* interaction with LIN28 than Class II family members. The difference is reproducible across multiple CLIP datasets independent of CLIP protocol modification (HITS-CLIP versus eCLIP), cellular contexts (HepG2, K562 cells and mESCs) and the targeted protein (LIN28A versus LIN28B). Importantly, the difference of the two subclasses in LIN28 binding affinity is clearly reflected in the complementary interactome capture assay (Treiber et al., 2017). This proposed model explains why isolated CSD does not bind, or binds only weakly, to human let-7a-1 (Nowak et al., 2017) and Xtr-pre-let-7f (Mayr et al., 2012), which do not have the GAU motif, but binds efficiently to let-7d, let-7f-1, and let-7g, which contain the GAU (Wang et al., 2017). Coincidentally, the crystal structure of LIN28 and let-7 was obtained for three Class I let-7 family members, but not any Class II family members (Nam et al., 2011). Similarly, in previous studies using LIN28 CLIP assays (Cho et al., 2012; Graf et al., 2013; Wilbert et al., 2012), the pre-let-7 members shown as examples for robust LIN28 binding are all from Class I.

**Figure 4:**
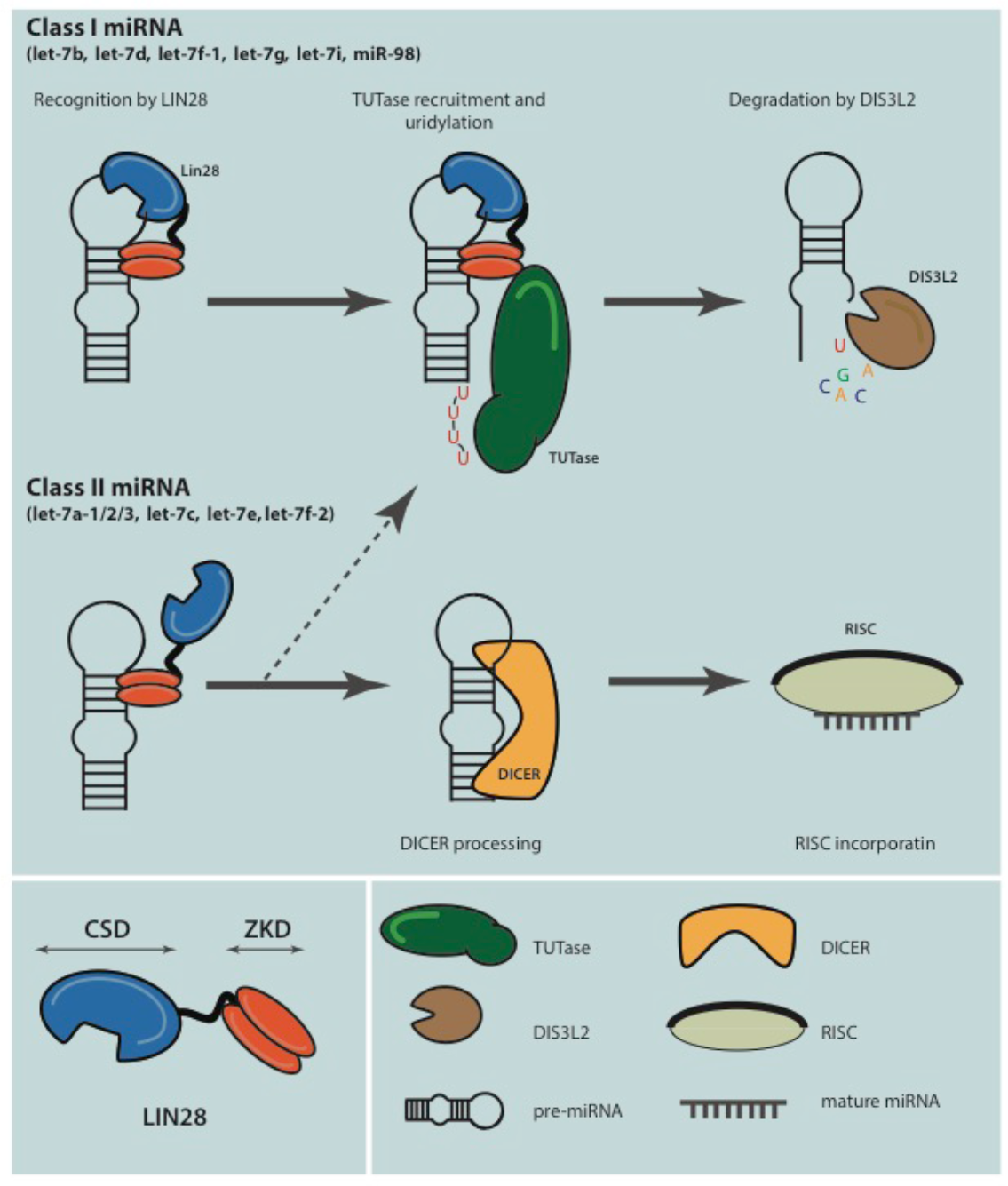
The proposed model of selective let-7 microRNA suppression modulated by the bipartite LIN28 binding. Related to Figure S6. Class I let-7 miRNA precursors have both CSD and ZKD binding elements, which efficiently recruit LIN28, leading to their 3′ uridylation by TUTase and degradation by DIS3L2. Class II let-7 miRNA precursors lack CSD binding element and are recognized by LIN28 with low efficiency, generally leading to an escape of these miRNAs from LIN28-mediated suppression.

The importance of the GAU motif for CSD binding helps to explain some unexpected observations reported in the literature. For example, to demonstrate the importance of the GGAG motif in the terminal loop region for LIN28-mediated uridylation and miRNA degradation, pre-mir-16-1, which lacks GGAG, was used as a negative control. However, relatively weak but reproducible binding of LIN28 to wild type pre-mir16-1 was observed *in vitro*, independent of mutations introduced in the ZKD that are expected to abolish its interaction with the GGAG motif (Heo et al., 2009). This interaction might be due to the presence of the GAU motif in the loop area of pre-mir-16-1. The GAU motif might also contribute to the observation that the chimeric pre-mir-16-1 with insertion of GGAG a few nucleotides downstream of the GAU element showed efficient LIN28 binding, 3′ uridylation and degradation, a point that was also previously noted (Nam et al., 2011).

The functional consequence of selective LIN28 binding to Class I versus Class II let-7 is clearly reflected in the much more efficient 3′ uridylation (observed from CLIP data of LIN28 and DIS3L2) and degradation (observed in HEK293 cells upon LIN28 overexpression or knockdown (Hafner et al., 2013; Wang et al., 2017; Wilbert et al., 2012)). Similarly, depletion of LIN28B in neuroblastoma cells using Cas9 targeting showed a much greater level of de-repression for Class I let-7 family members compared to Class II members (Powers et al., 2016). In these cancer cell lines, expression of let-7a members appears to be predominant among the let-7 family, despite high levels of LIN28 expression, suggesting that let-7a precursors lacking the GAU element are escaping LIN28 mediated repression (Hafner et al., 2013; Powers et al., 2016; Wilbert et al., 2012). However, we acknowledge that our proposed selective suppression model might not reconcile some discrepancies among previously reported results. For instance, several studies suggested that let-7a-1 (Class II) levels are low in ESCs and P19 cells, and that their abundance increases clearly upon LIN28A knockdown, indicating relatively efficient suppression by LIN28 (Heo et al., 2009; Viswanathan et al., 2008). However, the unbiased, quantitative comparison of Class I versus Class II let-7 members upon LIN28 knockdown in ESCs using deep sequencing remains lacking. Interestingly, we noticed that let-7a (generated from pre-let-7a-1/2/3; all Class II) and let-7f (generated from pre-let-7f-1 from Class I and pre-let-7f-2 from Class II) mature miRNAs are relatively abundant in ESCs compared to the other let-7 miRNAs expressed from the same poly-cistronic loci (Figure S6) and that let-7a is among the top 20 most abundant miRNAs (Wilbert et al., 2012), indicating that these Class II let-7 miRNAs might at least partially escape LIN28 suppression in ESCs. If the discrepancy is not due to technical differences, we cannot exclude the possibility that additional sequence or structural features (e.g., secondary RNA structures and other interacting proteins) in pre-let-7 can also affect LIN28-mediated suppression. Alternatively, there could be subtle differences between LIN28A and LIN28B in selectivity of Class I versus Class II let-7 pre-miRNAs as ESCs predominantly express LIN28A. So far, LIN28A and LIN28B are largely biochemically indistinguishable, although the two homologs were reported to have largely mutually exclusive expression in different cell lines and their subcellular localizations also differ, leading to potentially different mechanisms of action (Piskounova et al., 2011).

So what is the implication of this selective suppression model for developmental biology? On one hand, this model is consistent with the large number of let-7 family members in all bilateral animals, suggesting a strong evolutionary selection pressure to maintain diversity within the family. On the other hand, the benefit of having selective suppression remains a major, unanswered question. We propose several potential scenarios in which pluripotent stem cells might need to have a divergent subset of let-7 family members to escape LIN28 suppression. First, the selective suppression model could provide a mechanism for fine-tuning the abundance of let-7 expression. The timely and precise adjustment of the mature let-7 miRNA pool might be required to tightly control their downstream mRNA targets, including those essential regulators of stem cell pluripotency. Second, both LIN28A and LIN28B mRNAs possess let-7 binding sites on their 3′ UTRs and can themselves be subjected to suppression by let-7 (Rybak et al., 2008). In addition, let-7 also regulates the levels of MYC (Melton et al., 2010; Sampson et al., 2007), a transcriptional regulator of LIN28 (Chang et al., 2009; Dangi-Garimella et al., 2009). Such a complex multilayer regulatory feedback loop might be essential for the robust maintenance of the pluripotent state in ESCs, but will require a break during transition to the differentiated state. The subset of let-7 family members (Class II) that are capable of partially escaping from LIN28 suppression could provide such a trigger. Interestingly, during reprogramming of mouse embryonic fibroblasts towards induced pluripotent cells, greater efficiency was achieved by using let-7 antisense oligonucleotides compared to expression of LIN28 proteins, possibly due to the uniform suppression of all let-7 family members which cannot be achieved by LIN28 (Worringer et al., 2014). Finally, our analysis suggests that despite of their identical seed regions, each let-7 miRNA targets a unique, but not mutually exclusive, gene set (H.-S.C. and P.S., unpublished observation). Consequently, each target is potentially regulated by a set of Class I and Class II let-7 miRNAs, which act as selector switches to amplify the effects of fluctuations in LIN28 abundance. We hypothesize that the magnified response of Class I miRNAs to LIN28 upregulation leads to variable effects on the post-transcriptional regulation of let-7 targets, and that let-7 target expression profiles following LIN28 upregulation vary depending on the identities and classes of the regulating miRNAs. These possibilities do not have to be mutually exclusive, but they all suggest that the selective suppression model adds another layer to the remarkable complexity of the LIN28/let-7 axis not fully appreciated in previous studies.

## Acknowledgements

We thank members of the Zhang laboratory for helpful discussion. This study was supported by grants from the National Institutes of Health (NIH) (R01NS089676, R21NS098172 and R03HG009528 to C.Z.), the Simons Foundation Autism Research Initiative (307711 to C.Z.), and European Union’s Horizon 2020 Research and Innovation Programme (668858 to P.S.). High-performance computation was supported by NIH grants S10OD012351 and S10OD021764.

## Author contributions

DU and CZ conceived the study; DU, HSC and SMW performed data analysis; PS and CZ supervised the work; DU and CZ wrote the paper with input from all authors.

## Methods

### CLIP data processing

To determine the binding specificity of LIN28, we used Lin28a CLIP data derived from mouse embryonic stem cells (SRP012118) (Cho et al., 2012) and LIN28B eCLIP data derived from HepG2 and K562 human cell lines as part of the ENCODE project (https://www.encodeproject.org). For each dataset, raw reads were downloaded and processed using our established analysis pipeline CLIP Tool Kit (CTK) (Shah et al., 2017).

In analysis of the eCLIP data, slight modifications were made, as recommended by the original study. Specifically, the 3′ adaptors were trimmed using the cutadapt program (Martin, 2011), similar to the analysis pipeline used by the ENCODE consortium (--match-read-wildcards --times 1 -e 0.1 -O 1 --quality-cutoff 6 -m 18 -a $a1 -A ATTGCTTAGATCGGAAGAGCGTCGTGT -A ACAAGCCAGATCGGAAGAGCGTCGTGT -A AACTTGTAGATCGGAAGAGCGTCGTGT -A AGGACCAAGATCGGAAGAGCGTCGTGT -A ANNNNGGTCATAGATCGGAAGAGCGTCGTGT -A ANNNNACAGGAAGATCGGAAGAGCGTCGTGT -A ANNNNAAGCTGAGATCGGAAGAGCGTCGTGT -A ANNNNGTATCCAGATCGGAAGAGCGTCGTGT; $a1=NNNNNAGATCGGAAGAGCACACGTCTGAACTCCAGTCAC or NNNNNNNNNNAGATCGGAAGAGCACACGTCTGAACTCCAGTCAC, depending on the length of the degenerate barcode used for a specific library). After collapsing exact duplicates, the reads were subject to barcode removal and mapped to the reference genome (hg19) using bwa (Li and Durbin, 2009). Reads mapped to repetitive RNAs such as rRNAs and tRNAs as annotated in the RepeatMasker track were excluded. Potential PCR duplicates were further collapsed by modeling the random barcode to get unique tags. Only read2 (the read starting from 5′ end of the RNA tag) was used for analysis described in this paper. The unique tags were used for all downstream analysis, including visualization of read coverage in each genomic position.

To define LIN28 binding sites, replicates were combined to call CLIP tag clusters using a valley seeking algorithm (P≤0.05 after Bonforroni multiple testing correction; valley depth≥0.5). The sequences around the peak center (+/−100nt) were then extracted to evaluate the enrichment of the LIN28 consensus motif (GGAG and NGAU), using flanking sequences of the same size but 500 nt away from the peak center.

To define LIN28 binding sites at the single nucleotide resolution, we performed crosslink-induced mutation site (CIMS) analysis on the Lin28a CLIP data, as the protocol used to generate this dataset does not capture read-throughs at the crosslink sites. CIMS based on reproducible substitutions (FDR<0.05) were reported in this study. For the LIN28 eCLIP dataset, we performed crosslinking induced truncation site (CITS) analysis, as we observed minimal evidence of CIMS in this dataset. CITS with FDR<0.001 were reported in this study. Sequences around CIMS and CITS (−10,+10nt) were extracted for *de novo* motif analysis as described below.

### LIN28 *de novo* motif discovery

Currently, most of the software tools for de novo motif discovery (e.g., MEME (Bailey and Elkan, 1994) and HOMER (Heinz et al., 2010)) use a standard model with a position-specific weight matrix (PWM) to characterize the specificity of DNA- or RNA-binding proteins. Such a model is applied to a set of training sequences (e.g., sequences around CLIP tag peaks) to find the most over-represented sequence patterns allowing degeneracy. Since many RBPs recognize short and degenerate motifs, the reliability of this approach varies. To improve the precision of *de novo* motif discovery, we developed an algorithm which takes advantage of the single-nucleotide resolution map of protein-RNA interactions from CIMS and CITS analysis. This algorithm uses a model that augments the standard PWM model by jointly modeling RBP sequence specificity and the precise protein-RNA crosslink sites at specific motif positions at single-nucleotide resolution. As a result, this method reports both the sequence specificity of an RBP and the probability of crosslinking in each position of the motif. Details of the method will be described elsewhere. We used this algorithm to determine LIN28 binding motifs using CIMS from Lin28a CLIP in mESCs and CITS from LIN28B from human cell lines.

### Prediction of LIN28 binding sites in mRNA

To predict clusters of LIN28 motif sites genome-wide, we used our mCarts algorithm (Weyn-Vanhentenryck and Zhang, 2016; Zhang et al., 2013). We generated the positive training set from the significant peaks in the LIN28B eCLIP data from K562 cells, masking repeats, requiring a location within 1000 nt of an exon, and extending the peak center by 50 nt, resulting in 38,957 regions. The negative training set consisted of exonic regions extended by 1000 nt which did not overlap with any tags. mCarts was run to identify clusters containing at least 3 motifs, with motifs at most 30 nt apart. We generated three models: one searching for clusters of GGAGs, one searching for clusters of GATs, and one searching for clusters with any combination of GATs and GGAGs. These resulted in 214,152, 1,086,840, and 3,590,347 clusters, respectively. To evaluate the sensitivity of the results, we removed clusters overlapping with repetitive regions, ranked the clusters according to their score, and determined whether the cluster center overlapped with CLIP peaks (peak height region extended by 50 nt). We plotted the fraction of clusters overlapping CLIP peaks at each rank to compare the models.

### LIN28 structural visualization

All structural visualization of LIN28 and its targeted RNA let-7g was performed using PyMol software. All the data was retrieved from PDB (accession: 3TS2).

### LIN28 RNA-mediated protein pull-downusing let-7 pri-miRNA as a bait

To evaluate the binding affinity between different pre-let-7 family members with LIN28 and complement with CLIP data, we analyzed an interactome capture dataset using miRNA precursors as a bait (Treiber et al., 2017). Equal amount of each miRNA hairpin was used as a bait to capture associated proteins in 11 different cell lines. For each individual protein and each bait, the amount of capture was quantified by the spectrum counts from mass spectrometry analysis as the percentage total counts, averaged over all cell lines in which the protein was identified. The quantification used in our analysis was obtained from the original study.

### Uridylation analysis using ENCODE mock and LIN28B eCLIP data

The ENCODE project assayed over 100 RBPs using eCLIP in two human cell lines HepG2 and K562 (Van Nostrand et al., 2016), and data for each RBP consists of a mock and IP experiment. The mock experiment measures all captured RNA fragments crosslinked with any RBPs, so we estimated the expression levels of each miRNA by combining all generated mock experiments (94 in HepG2 and 92 in K562 at the time of this study) and counting the number of unique tags mapping to each pre-miRNA normalized by the total number of unique tags (read per million or RPM) in each sample. We estimated polyuridylation by identifying uridylated tags in LIN28B CLIP experiments (which should contain a stretch of Ts at the end of the read). To identify uridylated tags, we began with the unmapped LIN28B reads remaining after standard CLIP data processing. Using cutadapt (Martin, 2011), we first obtained the set of unmapped reads containing ≥4 consecutive Ts on the 3′ end and removed the Ts. These trimmed reads were then re-mapped to the genome and collapsed to identify unique uridylated tags using the CTK pipeline as described above. We then counted the number of uridylated reads on each miRNA precursor normalized by the total number of unique tags (expressed as RPM). Pre-let-7-g contains 3 T’s around the uridylation site (Ustianenko et al., 2016), so reads mapping to let-7-g were filtered to require ≥7 Ts. The coordinates of microRNA hairpins were based on miRBase R21 (June 2014) (Kozomara and Griffiths-Jones, 2014).

### Let-7 expression change upon LIN28 overexpression and knockdown

To evaluate the impact of LIN28 on let-7 expression, we used a miRNA-seq dataset from a published study (Hafner et al., 2013). This dataset was derived from HEK293 cells after expressing LIN28B for 72 hrs, after mock transfection (ctrl), and after LIN28B knockdown 72 hrs post-LIN28B siRNA transfection. The fold changes of let-7 expression from pairwise comparison were obtained from the original study.

### Selective suppression of let-7 by LIN28 in cancer

To investigate the correlation between LIN28 expression and let-7 expression we analyzed a panel of TCGA tumor samples of fourteen types in which LIN28A/B are sometimes reactivated. For each of these tumor types, primary tumors were profiled using both RNA-seq (Illumina Genome Analyzer or HiSeq RNA Sequencing Version 2) and miRNA-seq (Illumina HiSeq 2000 miRNA Sequencing) by TCGA. RNA-seq data that quantify mRNA expression level of 17,792 protein-coding genes, including LIN28A/B, were downloaded from the TCGA Data Portal (level 3 normalization; retrieved on 05/12/2015). We used log2(normalized count+1) in our analysis. miRNA expression estimates (level 3 normalization) were processed by Firehorse and downloaded from https://confluence.broadinstitute.org/display/GDAC/Dashboard-Stddata (Release: 2015_04_02 stddata Run). All the “NA” values were replaced by “0”. In our analysis, we used log2-transformed RPM (Reads Per Million miRNA mapped), and miRNA identities were taken from miRBase R21 (Kozomara and Griffiths-Jones, 2014).

To test whether let-7 miRNAs were enriched for correlation with LIN28, we performed gene set enrichment analysis of these miRNAs, in the context of all expressed miRNAs, as a function of miRNA correlation with LIN28A and LIN28B expression in tumors that showed LIN28A and LIN28B variability (median absolute deviation score > 0). GSEA (Subramanian et al., 2007) used weighted enrichment statistics and ratio of classes, with p-values computed using 1k gene-set permutations.

To evaluate the suppression of let-7 miRNAs by LIN28, we calculated the distance correlation (dCor) between LIN28A/B expression profiles and the profiles of each mature miRNA. We used dCor because of its ability to capture non-linear correlations (Szekely et al., 2007). Spearman’s correlation was used to determine the sign of dCor, which implies the direction of regulation. Only samples showing LIN28 presence, i.e., with nonzero read counts, were included for analysis. To estimate the significance of dCor, we shuffled the expression of LIN28A/B 1000 times and then calculated dCor between randomized LIN28A/B profiles and the profiles of all other protein-coding genes and mature miRNAs to produce nonparametric p-value estimates. We used a Mann-Whitney U test to compare distributions of distance correlations. We performed two types of comparisons: 1) We calculated dCor between the profiles of LIN28 and each miRNA species, including Class I and Class II let-7 miRNAs; then, we obtained the distribution and the average of dCor values within each subclass. 2) We summed up normalized expression across Class I let-7 miRNAs and Class II let-7 miRNAs (total expression) and calculated the dCor between total miRNA and LIN28 expression profiles.

